# Analysis of novel hyperosmotic shock response suggests “beads in liquid” cytosol structure

**DOI:** 10.1101/562728

**Authors:** A.I. Alexandrov, E.V. Grosfeld, A.A. Dergalev, V.V. Kushnirov, R.N. Chuprov-Netochin, P.A. Tyurin-Kuzmin, I.I. Kireev, M.D. Ter-Avanesyan, S.V. Leonov, М.O. Agaphonov

## Abstract

Proteins can aggregate in response to stresses, including hyperosmotic shock. Formation and disassembly of aggregates is a relatively slow process. We describe a novel instant response of the cell to hyperosmosis, during which chaperones and other proteins form numerous foci with properties uncharacteristic of classical aggregates. These foci appeared/disappeared seconds after shock onset/removal, in close correlation with cell volume changes. Genome-wide and targeted testing revealed chaperones, metabolic enzymes, P-body components and amyloidogenic proteins in the foci. Most of these proteins can form large assemblies and for some, the assembled state was pre-requisite for participation in foci. A genome-wide screen failed to identify genes whose absence prevented foci participation by Hsp70. Shapes of and interconnections between foci revealed by super-resolution microscopy indicated that the foci were compressed between other entities. Based on our findings, we propose a new model of the cytosol architecture as a collection of numerous of gel-like regions suspended in a liquid network. This network is reduced in volume in response to hyperosmosis and forms small pockets between the gel-like regions.

## Introduction

Systems biology aspires to understand life in quantitative detail, eventually allowing complete modeling of living cells *in silico*. For such an endeavor, it is crucial to predict distribution and local concentrations of substances and molecular assemblies within a cell under varying conditions. However, even for simple cytosolic processes, this has been difficult to achieve due to various aspects of cytoplasmic heterogeneity (for a review, see (Luby-Phelps, 1999; Luby-Phelps, 2013) and differing views on the nature of the cytosol, i.e. is it a simple solution, crowded liquid, or a hydrogel (Grygorczyk et al., 2015). While all of these models describe some properties of the cytosol, they are difficult to unite in a single framework and thus there is a lack of comprehensible mechanistic models. Notably, the cytoskeleton, which accomplishes most of the active transport in the cell, has been implicated in cytosolic structuring (Hu et al., 2017; Provance et al., 1993), however, it does not seem to fully account for existing observations (Weiss et al., 2004). Another difficulty that adds to the complexity of cytosolic structure is the ability of the cytosol to change its viscosity in a dramatic manner, such as during changes of pH (Munder et al., 2016; Parry et al., 2014) or osmotic pressure (Miermont et al., 2013).

Hyperosmotic shock is a ubiquitous environmental factor commonly encountered by microorganisms and multicellular organisms. It is also relevant for some tissues in mammals (Brocker et al., 2012). The response of cells to hyperosmosis has been studied in great detail in terms of the sensing of hyperosmotic shock, the signaling cascades which mediate the cells’ responses, and the mechanisms of adaptation to hyperosmotic conditions via synthesis and retention of osmolytes, namely glycerol, in yeast (reviewed in (Saito and Posas, 2012)).

However, the immediate consequences of hyperosmotic shock are less well characterized.

During hyperosmotic shock cells shrink due to water efflux. This is accompanied by increased cytoplasmic viscosity and reduction of diffusion rates for various proteins (Miermont et al., 2013), as well as aggregation of cellular proteins and model amyloidogenic proteins in *C. elegans* (Burkewitz et al., 2011) and yeast (Han and Emr, 2011; Oeser et al., 2016). Also, in yeast, hyperosmotic shock can influence the disappearance and appearance of prion amyloids (Newnam et al., 2011; Tyedmers et al., 2008). Notably, protein aggregation as well as formation of visible foci in response to various stresses takes a noticeable amount of time, i.e. at least several minutes for severe heat shock. Dissolution of aggregates and foci is an even longer process which can take up one or more hours (Wallace et al., 2015).

Another class of entities that can appear in response to changing conditions are protein droplets that form due to liquid-liquid phase separation (LLPS) (reviewed in (Hyman et al., 2014; Shin and Brangwynne, 2017)). LLPS can proceed in a matter of seconds in some cases, for instance, when proteins prone to aggregation are rapidly brought into close proximity (Bracha et al., 2018b). Notably, a recent study reported that hyperosmosis caused formation of phase-separated droplets in mammalian cells (Cai et al., 2018).

Gathering of a particular protein into aggregates or other types of assemblies can be monitored *in vivo* if it is labeled with a fluorescent moiety e.g. green fluorescent protein (GFP). Here we observed that some proteins fused to GFP rapidly formed reversible intracellular foci in response to hyperosmotic shock (OSF, **O**smotic **s**hock **f**oci). The shape and dynamics of appearance and disappearance of OSFs were inconsistent with classic protein aggregation, but indicated formation of highly reversible entities with unusual properties and morphology. This led us to propose a new model of the cytosol as gel-like “beads” suspended in a liquid network.

## Results

### Сhaperones can form cytoplasmic foci in response to hyperosmosis

Earlier it was observed that hyperosmotic shock causes aggregation of cellular proteins and model amyloidogenic proteins in *C. elegans* (Burkewitz et al., 2011), as well as influences disappearance and appearance of prion amyloids in yeast (Newnam et al., 2011; Tyedmers et al., 2008). Since some chaperone proteins were shown to bind to aggregates of misfolded proteins, one could expect that GFP fusions of such chaperones would decorate aggregates formed in response to hyperosmotic shock, thus allowing monitoring of aggregate formation *in vivo*. To do this, the strains from the Yeast GFP fusion collection (Huh et al., 2003) containing tagged Hsp104 and Ssa1 chaperones, which bind both to amorphous and amyloid aggregates (Chernoff et al., 1995; Glover and Lindquist, 1998) and are expressed at relatively high levels, were initially used. In order to study protein aggregation in response to hyperosmotic shock, we monitored the localization of these proteins, as well as several other tested chaperones. We observed that they formed numerous foci, which we term OSFs (Figure 1), in response to various types of hyperosmotic stress (high concentrations of KCl, sorbitol and glycerol in culture medium). Further experiments with Ssa1, unless noted otherwise, were performed using the Ssa1-Dendra2 (Ssa1-DDR) fusion protein, which behaved identically to Ssa1-GFP in terms of OSF formation, but did not form single large and stable inclusions similar to those observed in (Kaganovich et al., 2008), which complicated OSF visualization. Also, KCl was used as the hyperosmotic agent in further experiments.

**Figure 1.**
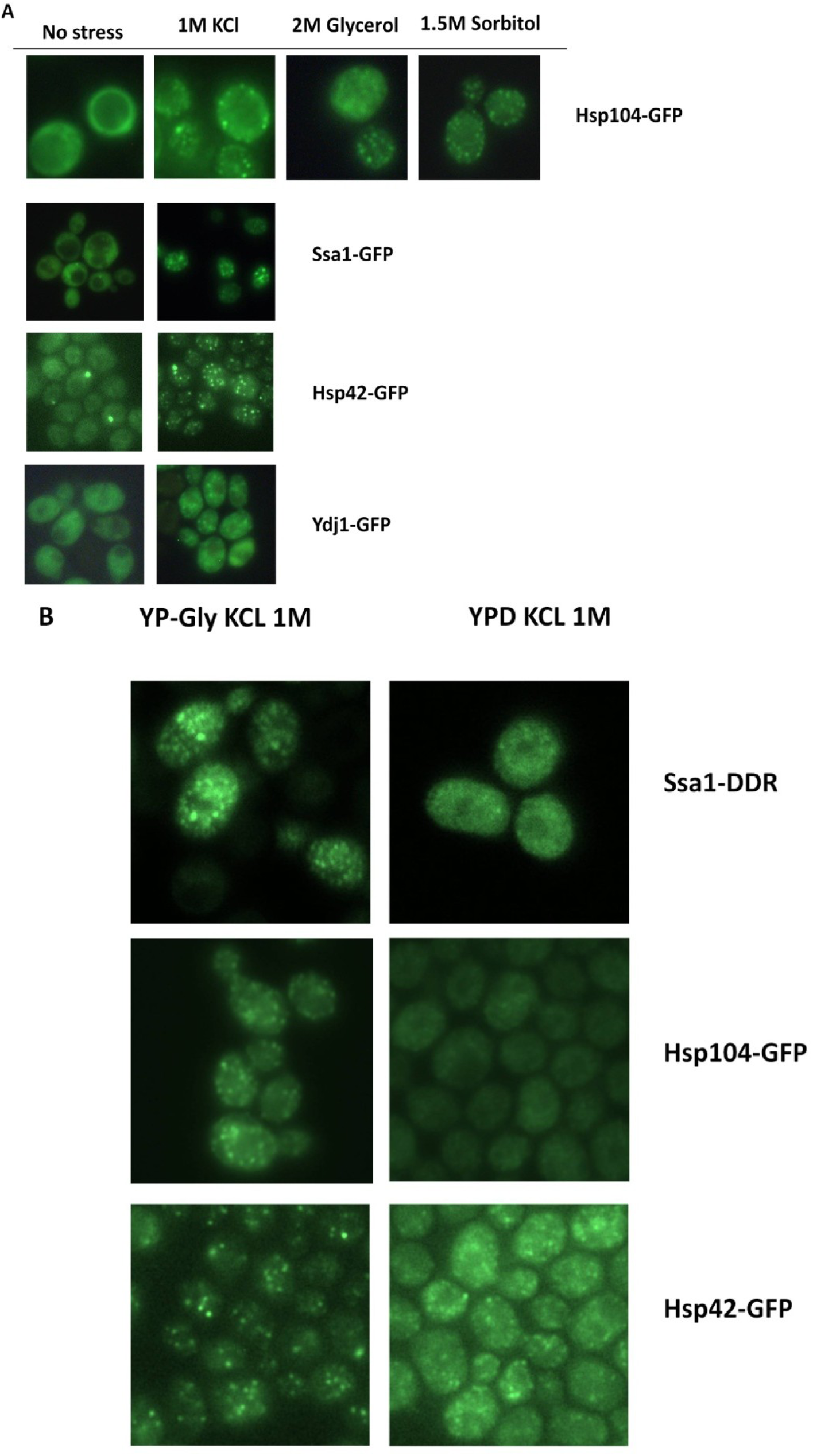
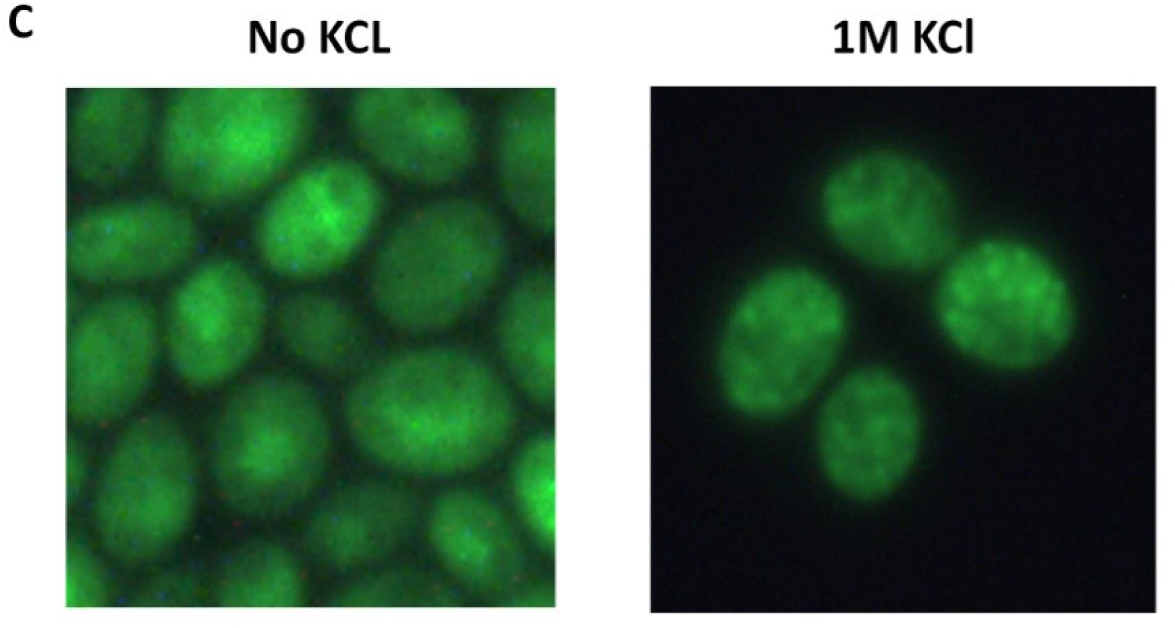
Сhaperones can form OSF under various stress conditions both in *S. cerevisiae* and *O. parapolymorpha*. **(A)** Cells of the BY4741 strain bearing the indicated GFP-fusion protein were grown in YP-Gly medium and then transferred onto medium with the indicated concentration of KCl, glycerol or sorbitol. **(B)** Cells bearing the indicated GFP-fusion chaperone were grown in YP-Gly or YPD medium to logarithmic phase and transferred onto the same medium supplemented with 1M KCl **(C)** Cells of O. parapolymorpha, producing the closest Ssa1 homologue tagged with GFP (see materials and methods) were grown in YP-Gly medium to logarithmic phase and transferred onto YPD medium with 1M KCl.

Ssa1-DDR formed OSFs efficiently when cells were grown to high density in YPD medium, while cells of logarithmic cultures were not capable of forming OSFs or formed indistinct OSFs. This suggested that OSF formation by Ssa1 was dependent on the carbon source, since upon reaching a certain density a yeast culture depletes glucose in the medium by converting it to ethanol, at which point the culture experiences the so-called diauxic shift, and starts consuming ethanol generated via glucose fermentation. In agreement with this suggestion, cells producing Ssa1-DDR and Hsp104-GFP on medium with glycerol as the sole carbon source formed OSFs efficiently even in logarithmic cultures (Figure 1B). On the other hand, the chaperone Hsp42 formed distinct OSFs not only in cells grown on non-fermentable carbon source such as glycerol, but also in cells consuming glucose (Fig. 1B). This indicates that OSF formation by different proteins may be accomplished under different conditions.

To understand if OSF formation is conserved in other yeast species, we assayed OSF formation by Hsp70 in the distantly-related methylotrophic yeast *Ogataea parapolymorpha*. Microscopic analysis of cells expressing the closest homologue of *S. cerevisiae* Ssa1 fused to GFP demonstrated that this chimeric protein also formed OSFs (Figure 1C).

To observe OSFs at increased resolution, we used Structural Illumination Microscopy (Gustafsson, 2000; Heintzmann and Cremer, 1999) and subsequent image deconvolution. Interconnections between some of the OSFs became evident, while others exhibited small protuberances which did not contact other foci (Figure 2). Overall, the OSFs seemed to form a more or less interconnected network in the cell.

**Figure 2.**
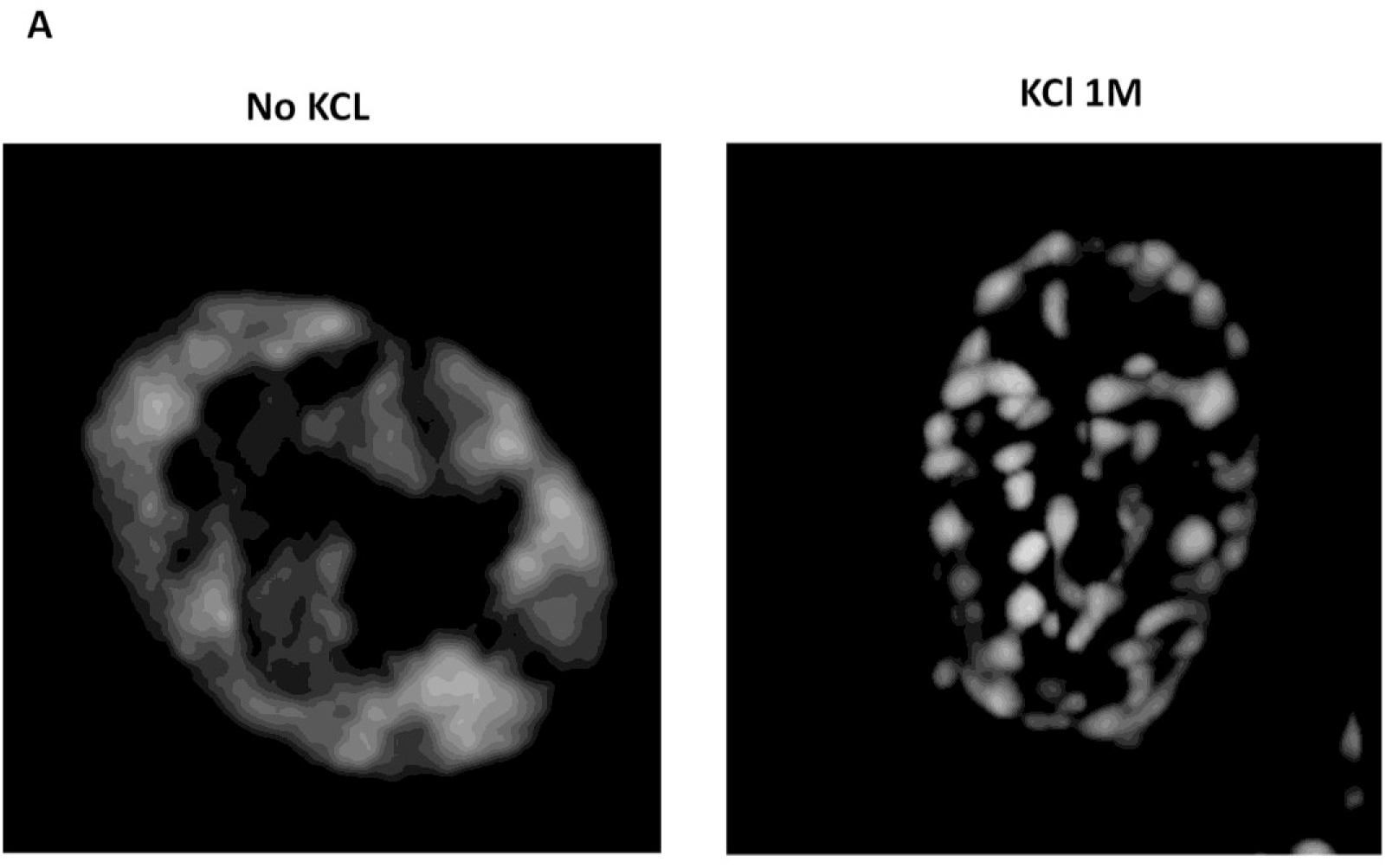
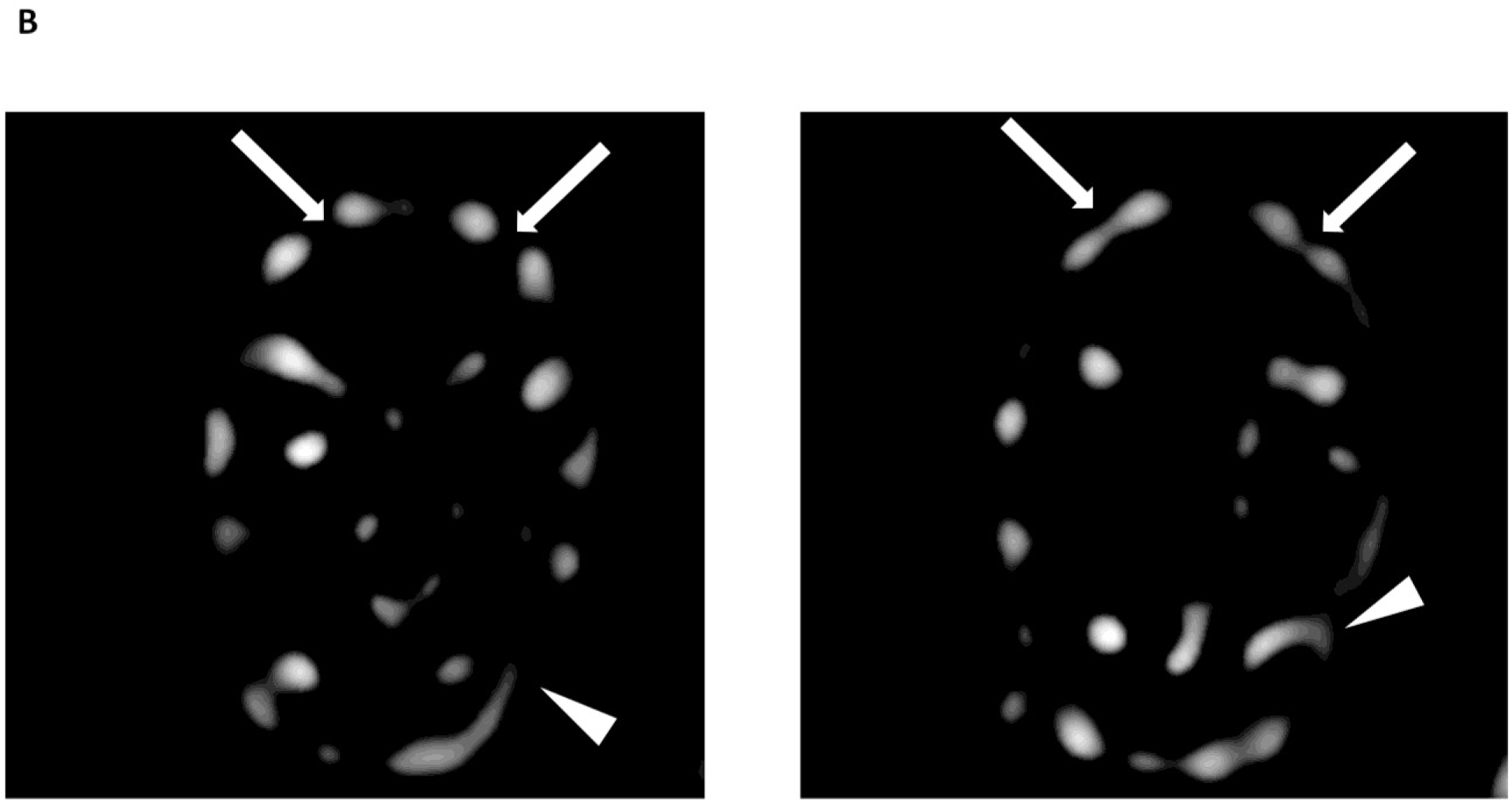
Structured Illumination Microscopy reveal bridges between OSFs as well as small OSF protuberances. Cells producing the Ssa1-DDR protein were collected in the logarithmic phase of growth from YP-Gly medium, and either subjected or not subjected to hyperosmotic shock with 1M KCl in the same medium. **(A)** Maximum intensity projection of a representative shocked and non-shocked cell. (**B)** Selected optic sections from the same shocked cell shown in (A) which contain OSFs with protuberances or interconnections between adjacent OSFs. Arrows indicate interconnections while protuberances are indicated by triangles.

### OSF formation is rapid and reversible

To determine how fast OSFs form and whether they persist after stress removal, we performed time-lapse microscopy of cells exposed to sequential hyperosmotic and normoosmotic conditions. Cells grown on normosmotic medium were imaged in real time during addition of hyperosmotic agent. Then the hyperosmotic medium was removed and replaced with normosmotic medium. To our surprise, OSF formation was very rapid, i.e. the cells formed OSFs within seconds after exposure to KCl and these OSFs disappeared just as rapidly upon cessation of hyperosmosis (Figure 3 and video). This makes it highly unlikely that OSFs represent protein aggregates or stable protein complexes. Next, we used image analysis to estimate the volume of observed cells. We showed that OSFs appear during shrinking after addition of KCl and that, upon shock cessation, the foci disappear concomitantly with сytoplasmic volume restoration (Figure 3).

**Figure 3.**
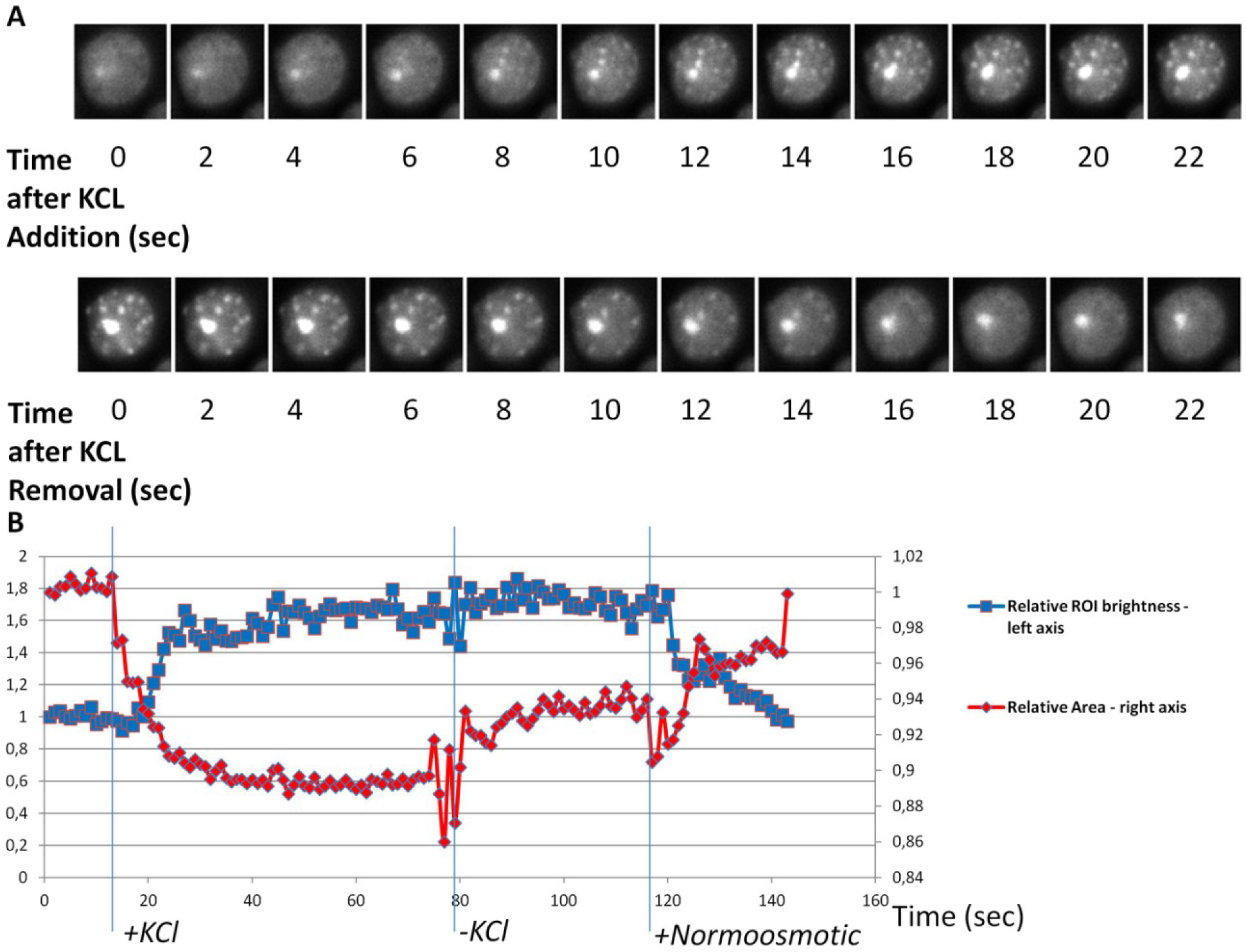
Dynamics of Ssa1-GFP OSF formation and disappearance. **(A)** Timelapse images of OSF formation and disappearance in response to onset and removal of hyperosmosis, respectively, obtained using cells expressing the Ssa1-GFP protein. Images were obtained using confocal real time microscopy. **(B)** Timecourse of changes in cytoplasmic area and OSF formation/disappearance. Vertical lines depict the timepoints at which the KCl-induced hyperosmotic shock was administered (+KCl), the time at which medium containing KCl was aspirated (-KCl), as well as the time at which normoosmotic medium was added (+Normoosmotic). The presented graphs were obtained from the cell images in **(A)**, except with higher temporal resolution. 10 other cells were also used to construct graphs with similar results. The Ssa1-GFP protein was used instead of Ssa1-DDR to provide stronger signal without photobleaching. The large aggregate that is constant through both timecourses is an IPOD-like inclusion of Ssa1-GFP that has no relation to OSFs. Fluctuation of the graphs during KCl removal and slight change of apparent cell area after it is due to vibrations of the sample during solution aspiration, which affected microscope focusing. –KCl неясно

### Identification of OSF-forming proteins by screening strains of the GFP-fusion library

Thus, the rapid and reversible formation of OSFs, coupled to their unusual shape, indicates that these structures are likely to be of a soft, probably liquid, nature. Most likely, due to these reasons, we failed to isolate OSFs of Ssa1-DDR by centrifugation of lysates of cells (data not shown), even if cell lysates were obtained under increased crowding conditions, which has been shown to facilitate purification of unstable protein complexes (Petrovska *et al*., 2014). Therefore, in order to identify proteins capable of OSF formation, we used high-throughput microscopy to screen for changes in protein localization in response to hyperosmotic shock in a collection of 4156 strains expressing GFP-fusion proteins (Huh *et al*. 2003). Cells were grown in YP-Gly medium, washed, and then resuspended in 1M KCl or distilled water, transferred to 384-well glass bottom microscop**y** plates and analyzed by high-throughput microscopy. This approach restricted adjusting exposure time in each strain, so we chose to use the same exposure time over the screening, which could be inappropriate for observation of a number of poorly expressed proteins. Despite these limitations, 10 novel OSF forming proteins were revealed (Table 1 and Supplemental material). These included proteins of unknown function, metabolic enzymes, a subunit of the translation initiation factor (Clu1) and several chaperones. Notably, neither the control protein Dendra2 expressed from a strong *ADH1* promoter, nor most of the tested GFP fusion proteins, including highly expressed ones, such as Pab1-GFP, Pub1-GFP and Tdh1-GFP formed OSFs (Figure 4). Importantly, analysis of cells expressing pairs of OSF-forming proteins and growing in conditions promoting OSF formation showed that they could colocalize (Figure 5). This was shown for pairs of chaperones that are known to interact with each other, i.e. Hsp104 - Ssa2 and Ydj1 – Ssa2, as well as for pairs of functionally unrelated proteins, such as Gln1 - Ssa2 and Tps1 – Hsp104.

**Figure 4.**
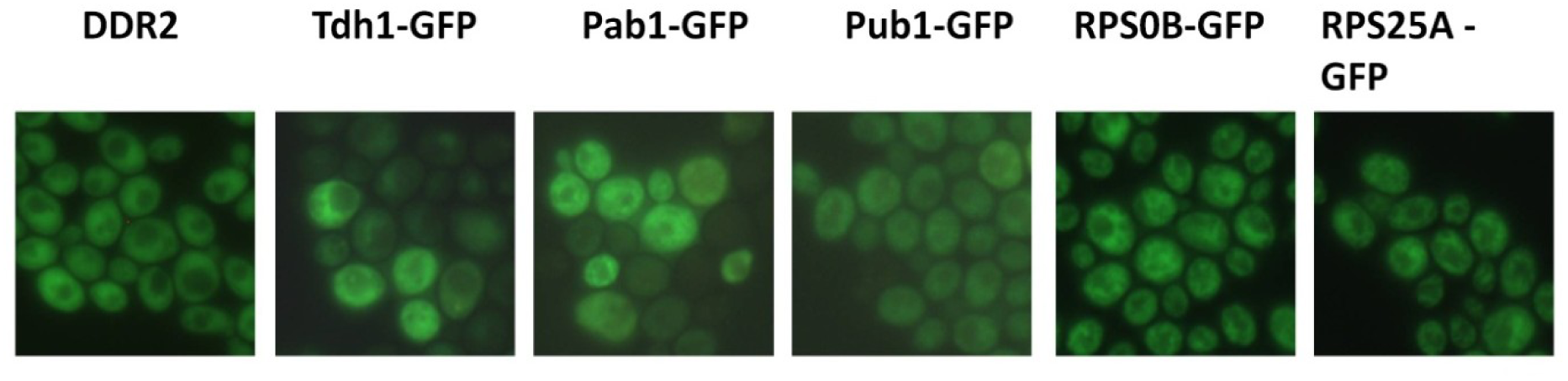
Examples of cytoplasmic proteins that do not form OSFs. *Cells producing the indicated GFP fusion proteins were grown on YP-Gly medium to log phase and transferred onto the same medium with 1M KCl.*

**Figure 5.**
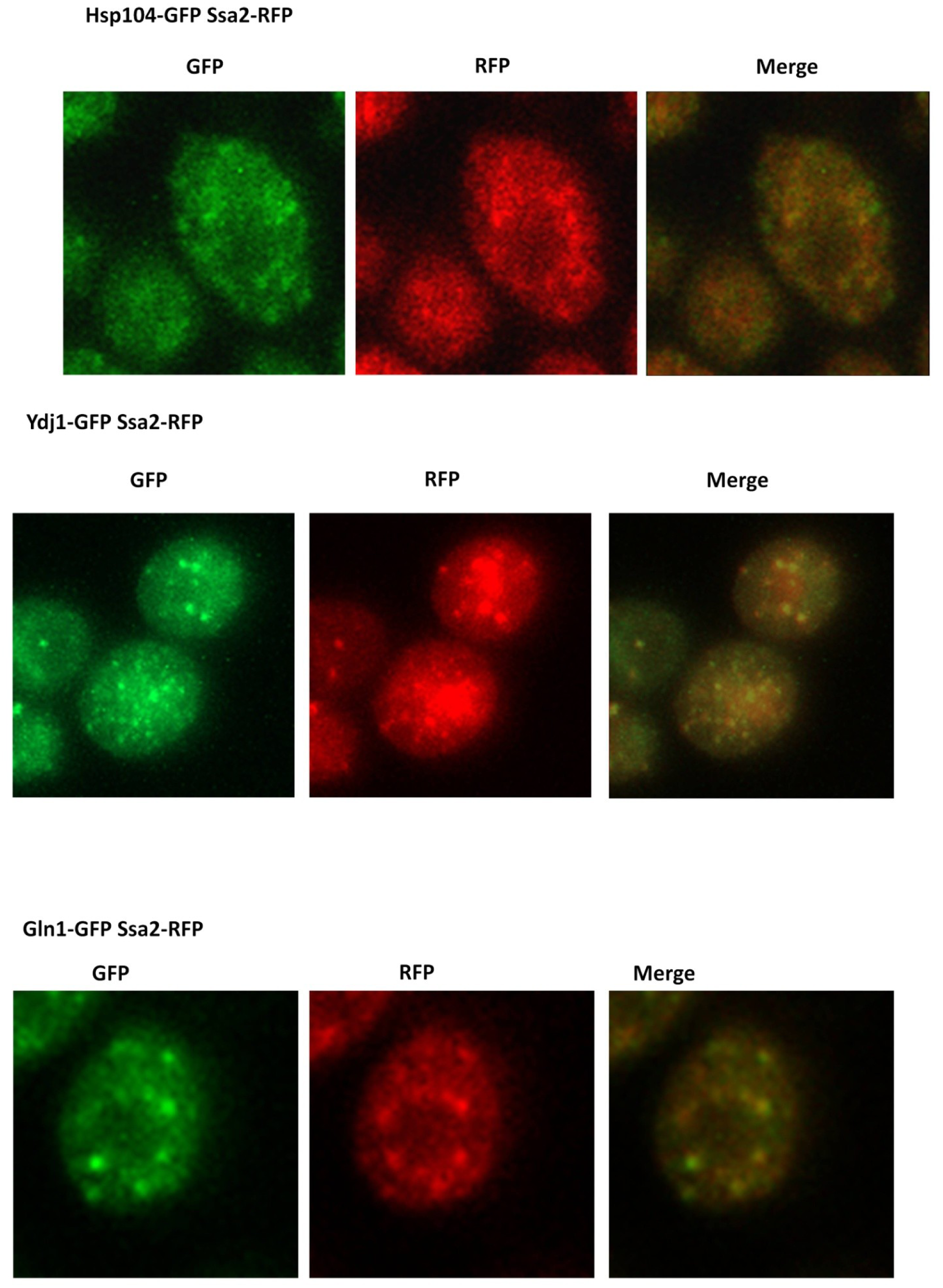
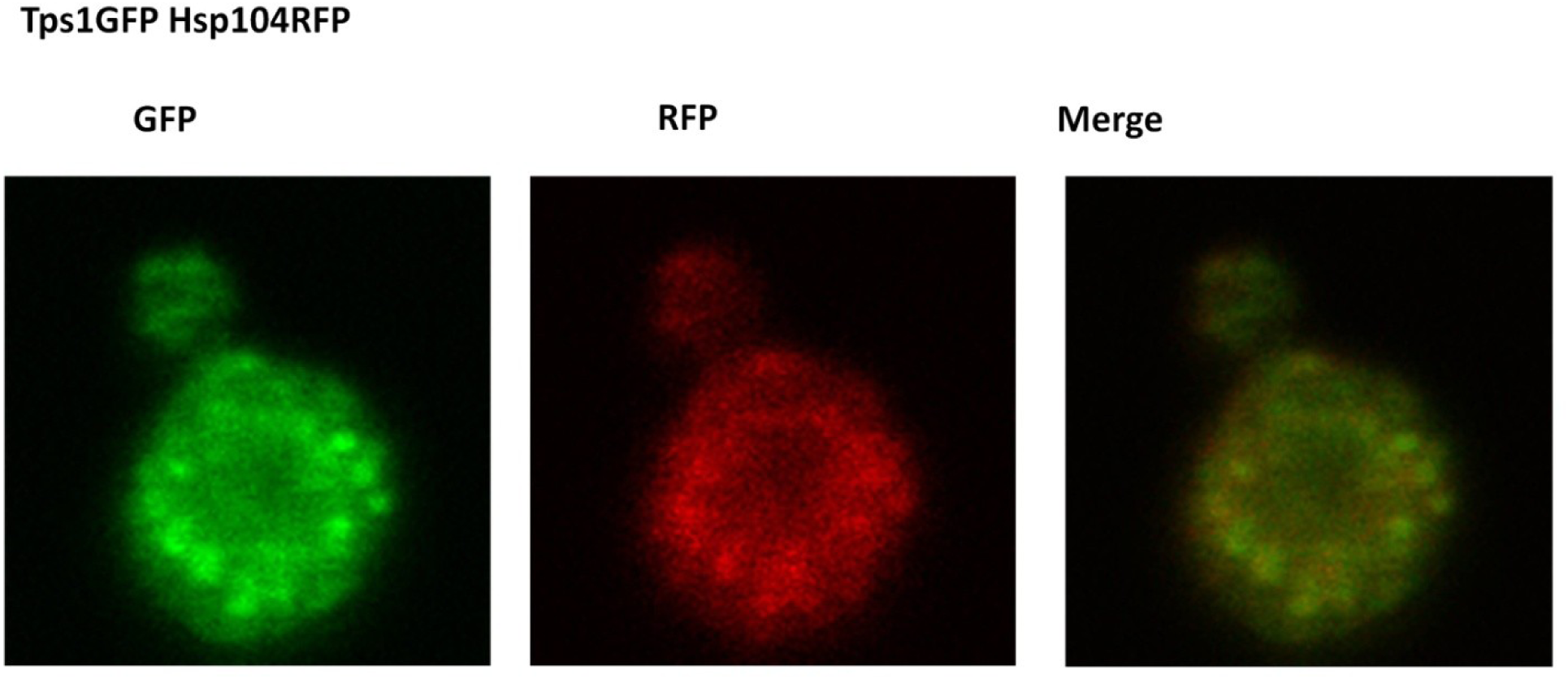
Proteins colocalize to the same OSFs during hyperosmotic shock. *Cells expressing pairs of the indicated GFP-tagged proteins were grown to mid-log phase in YP-Gly medium, transferred onto the same medium with 1M KCl and visualized by confocal fluorescent microscopy.*

**Table 1.** OSF-forming proteins identified in this work.

|  | **Protein** | **Function** | **Identified via** | **Large assembly formation, reference** |
| --- | --- | --- | --- | --- |
| 1 | Hsp104 | Chaperone | Designed choice | + |
| 2 | Ssa1 | Chaperone | Designed choice | + |
| 3 | Ssa2 | Chaperone | Designed choice | + |
| 4 | Ydj1 | Chaperone | Designed choice | + |
| 5 | Sis1 | Chaperone | GFP-fusion screening | + |
| 6 | Hsp42 | Chaperone | GFP-fusion screening | + |
| 7 | Hsp26 | Chaperone | GFP-fusion screening | + |
| 8 | Tsa1 | Thioredoxin peroxidase, targets Hsp70 to oxidation-damaged proteins | GFP-fusion screening | +<br>(Hanzén et al., 2016) |
| 9 | Tps1 | Trehalose synthesis during severe stress | GFP-fusion screening | +<br>(Londesborough and Vuorio, 1991) |
| 10 | Fas2 | Fatty acid synthase | GFP-fusion screening | + (Lomakin et al., 2007) |
| 11 | YGL082W | Putative protein of unknown function; predicted prenylation/proteolysis target of Afc1p and Rce1p | GFP-fusion screening | ? |
| 12 | Clu1 | Subunit of the eukaryotic translation initiation factor 3 (eIF3) | GFP-fusion screening | possibly +, component of large eIF3 complex (Vornlocher et al., 1999), interacts with mRNA |
| 13 | Ape1 | Vacuolar aminopeptidase | GFP-fusion screening | + (Kim et al., 1997) |
| 14 | Gln1 | Glutamine synthetase | GFP-fusion screening | + (He et al., 2009; Petrovska et al., 2014) |
| 15 | Dcp2 | P-body component, RNA-decapping | Designed choice | probably +, when interacting with mRNA |
| 16 | Edc3 | P-body component, RNA-decapping | Designed choice | probably +, when interacting with mRNA |
| 17 | PrD-Mot3 | Transcriptional repressor, prion protein | Designed choice | + (Alberti et al., 2009) |
| 18 | PrD-Pan1 | Endocytic protein with prion-like domain | Designed choice | + (Alberti et al., 2009) |
| 19 | Prd-Sup35 | Translation termination factor, prion protein | Designed choice | + (Alberti et al., 2009; Wickner et al., 1995) |

### Identification of additional OSF-forming proteins

Earlier it was described that upon hyperosmotic shock some of the stress-granule and processing-body (P-body) proteins can form foci which are assumed to be P-bodies (Huch and Nissan, 2017; Ramachandran et al., 2011; Teixeira et al., 2005). In agreement with this we observed that the P-body marker proteins Dcp2 and Edc3 tagged with GFP formed foci (Figure 6A), though stress-granule markers Pub1 and Pab1 did not (Figure 5). Even though the P-body proteins are not highly expressed, we still managed to detect OSF-like foci after approximately 30 sec of exposure to 1M KCl (the minimal post-shock time we could achieve). Notably, previous work on P-body formation in hyperosmotic conditions did not use such short shock exposure times. Importantly, the dynamics of OSF formation/disappearance by Dcp2-GFP and Edc3-GFP was so rapid, that they are difficult to equate to the much more stable P-bodies that appear in response of glucose deprivation in culture medium (Teixeira et al., 2005), thus indicating that the P-body protein foci that form during hyperosmotic shock may not be genuine P-bodies. More specifically, we suggest that some large, but microscopically unobservable complexes related to mRNA-processing may be constantly present in the cell, and hyperosmotic shock causes concentration of these complexes (see discussion).

**Figure 6.**
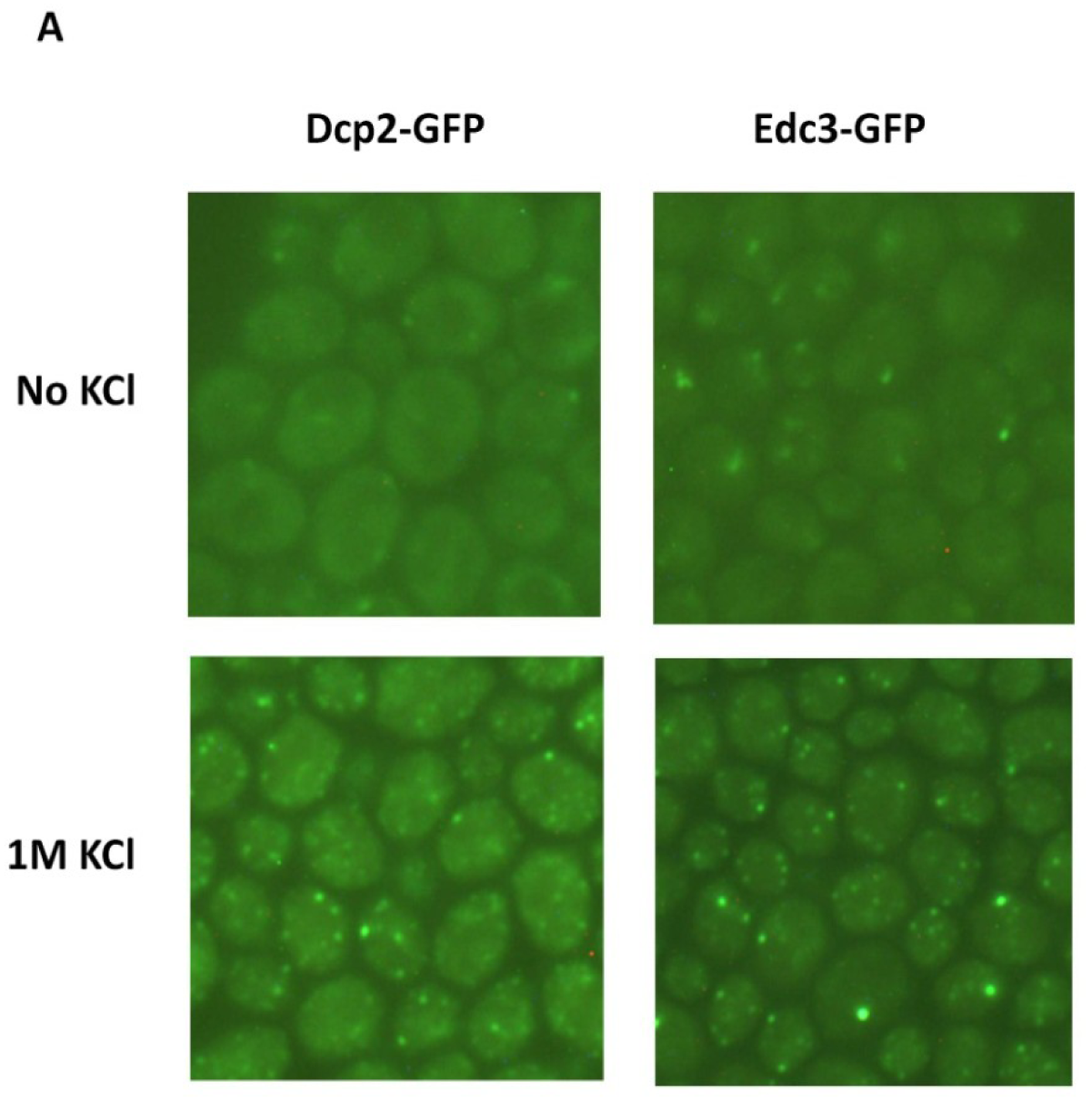
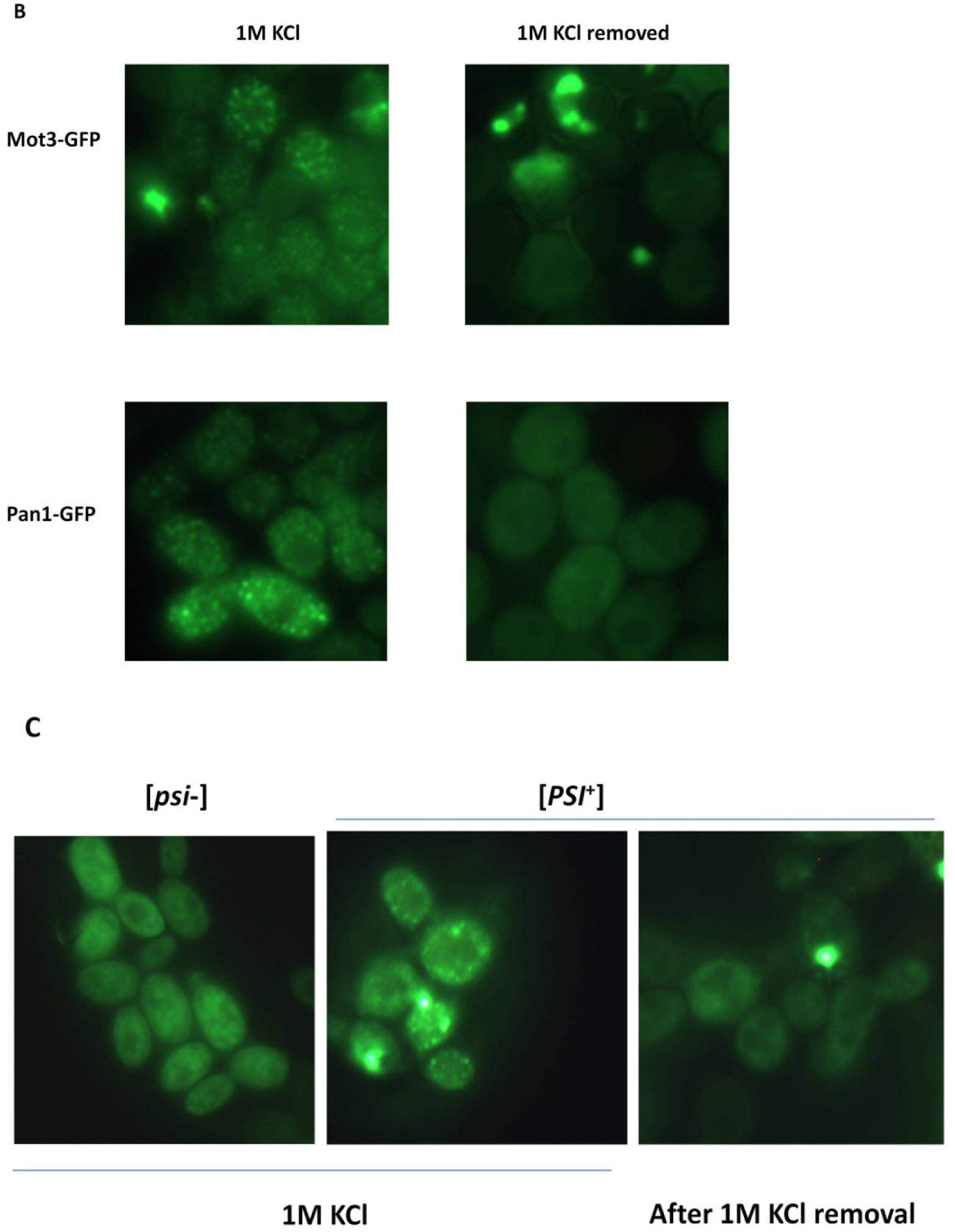
P-body proteins and proteins with amyloidogenic domains can form OSFs. **(A)** *Cells producing the indicated GFP-fusion proteins, were grown in YP-Gly medium, and washed in SC-Gly medium to reduce background medium fluorescence. The cells were subsequently subjected to hyperosmotic shock (1M KCl) in SC-Gly **(B)** Cells of the 74D-694 [psi^-^] strain harboring the plasmids expressing PrDMot3-GFP or PrDPan1-GFP were grown in SC-Gal medium to induce GFP-fusion protein production and then transferred onto the same medium supplemented with 1 M KCl. **(C)** Cells of the 74D-694 strain harboring the plasmid expressing PrDSup35-GFP, as well as containing or lacking prion amyloids of Sup35 ([PSI^+^] or [psi^-^], respectively) were grown in SCGly medium and then transferred onto the same medium with 1 M KCl. The growth of cells in SCGly was required to observe diffuse fluorescence of the protein in normoosmotic conditions, since high production of PrdSup35-GFP upon growth of [PSI^+^] cells in SC-Gal resulted in almost complete accumulation of this protein in IPOD-like inclusions. Due to the presence of the ade1-14 mutation, additional adenine was added to the medium to reduce of the amount of autofluorescent red pigment accumulated in cells of the ade1 mutants.*

**Figure 7.**
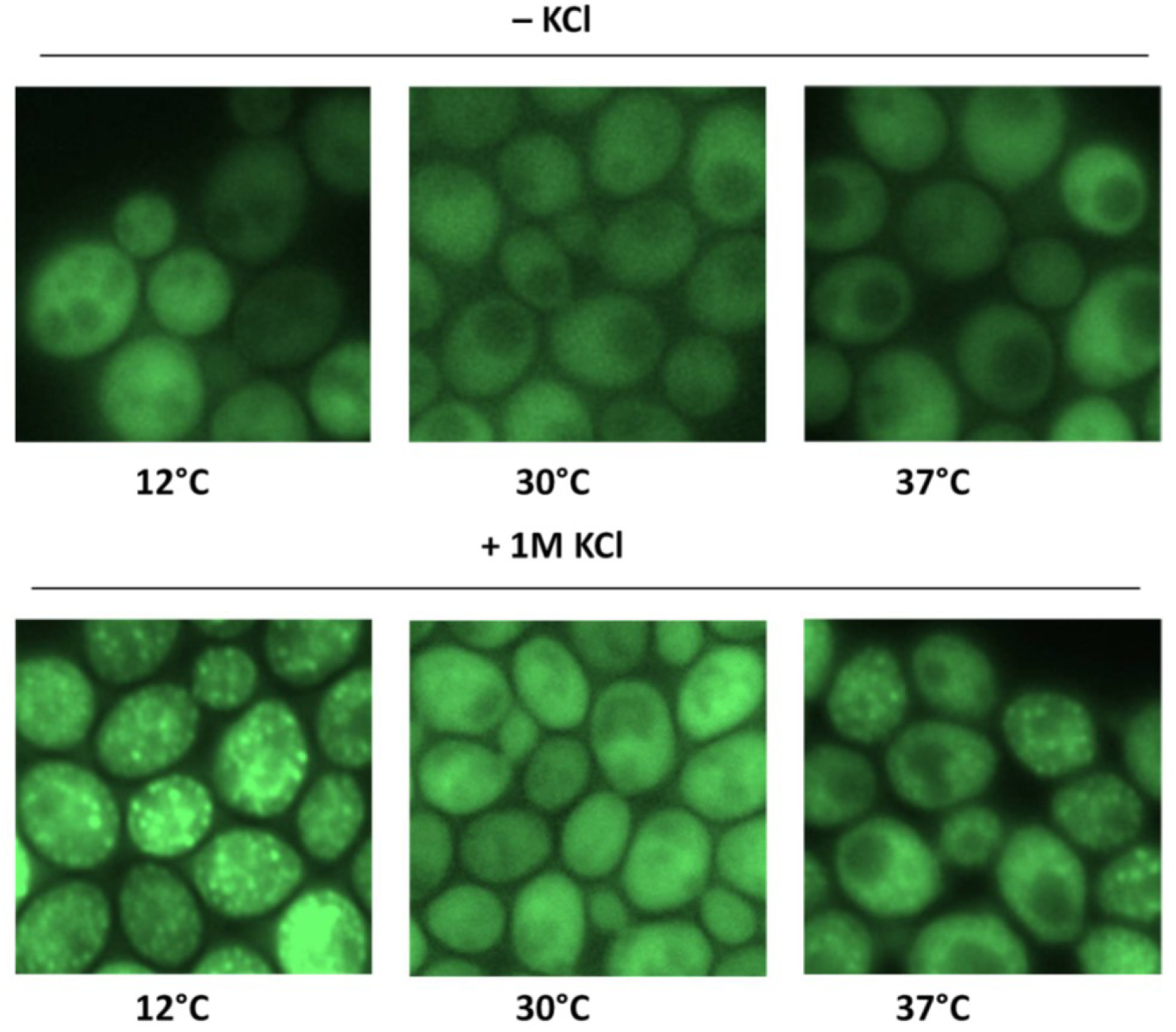
Conditions of mild stress during growth stimulate formation of OSFs by Ssa1-DDR. *Cells of the BY4741 strain bearing the Ssa1-DDR fusion protein were grown in YPD under the indicated conditions and then subjected to hyperosmotic shock. Exposition times were adjusted to have approximately equal intensity in the various samples.*

Formation of numerous small foci in response to hyperosmosis has been reported for the interacting transcriptional regulators Tup1 and Сyc8 (Han and Emr, 2011; Oeser *et al*., 2016). Notably, formation of foci for a mutant form of Cyc8 was dependent on its Q/N-rich domain (Oeser et al., 2016), which is required for amyloid formation (Patel et al., 2009). To determine whether other amyloidogenic proteins could form OSFs, we assayed OSF formation by overproduced prion and prion-like domains (PrD) of three such proteins fused to GFP, namely PrDSup35-GFP, PrDMot3-GFP and PrDPan1-GFP. These proteins were capable of forming OSFs similar to those of Ssa1-DDR OSFs in terms of their rapid formation and disassembly (Figure 6B). The rapid disassembly of these OSFs was especially surprising, since it indicated that even though these proteins can form amyloid aggregates upon overproduction (Alberti *et al*., 2009), the OSFs which they formed were not stable.

### Large complex formation is required for formation of OSFs by some proteins

Notably, PrD-Sup35-GFP was able to form OSFs only in the cells possessing prion determinant [*PSI+*], which implies involvement of this protein in prion polymers (Figure 6C). This suggested that OSFs are formed only by proteins that are members of large complexes, while monomeric proteins do not exhibit such behavior.

Perusal of the data on the identified proteins showed that nearly all of the identified proteins could form large protein assemblies (See Table1 and Discussion) in the megadalton range. The OSF-forming proteins that we had identified were especially enriched in chaperone proteins, and we had already observed that growth on a non-fermentable carbon source stimulated OSF formation by Ssa1. Such an environment is known to increase levels of oxidatively damaged proteins (Vasylkovska et al., 2015), so we reasoned that for chaperones, stressful conditions might increase the amount of chaperone bound to damaged proteins and thus increase the share of chaperone involved in the formation of large protein assemblies, thus stimulating the ability to form OSFs.

To test this, we assayed how heat stress (37°C) and cold stress (12°C) affected OSF formation and discovered that these treatments stimulated formation of OSF, while other treatments such as ER-stress did not (data not shown).

### Screening of the yeast gene knock-out/down strain libraries for genes essential for OSF formation

To assay the genetic control of OSF formation, we used the Synthetic Genetic Array (SGA) technique (Tong, 2001) to introduce Ssa1-DDR into strains of the commercially available Yeast Deletion (Giaever et al., 2002) and Yeast DAmP collections (Breslow et al., 2008). We then used the resulting array of strains to search for gene deletions or knock-downs which prevent formation of Ssa1-DDR OSFs in hyperosmotic conditions. Approximately 6000 strains were grown on YP-Gly plates, transferred to 1M KCl and then subjected to high-throughput microscopy. After obtaining preliminary results and rechecking the observed phenotypes of strains, we could not identify any genes which reproducibly prevented OSF formation.

## Discussion

In this work we discovered a novel type of cellular response to hyperosmotic shock – the nearly instant appearance of foci formed by GFP-labeled proteins and identify a number of proteins that form these OSFs. Intriguingly, the fast dynamics of OSF formation and disassembly, as well as tight correlation of their appearance and disappearance with changes of the cytoplasmic volume make it unlikely that OSFs represent conventional protein aggregates or stable protein complexes. This conclusion is also supported by the observation that even though Sup35 and Mot3 proteins are capable of forming highly stable amyloid aggregates (Alberti et al., 2009), they form labile OSFs which are disassembled in a matter of seconds after shock removal.

Of the available models which could explain our data, the process of liquid-liquid phase separation, which has attracted much interest in the last few years, might be relevant. In frame of this process, OSFs could be liquid droplets suspended in the cytosol, which form rapidly due to any of the following factors that accompany hyperosmotic shock – increased protein concentration, increased crowding, increased ion concentrations. Notably, a recent preprint on BioRXiv reports foci similar to OSFs formed by the YAP protein in response to hyperosmosis in mammalian cells (Cai et al., 2018). The authors base their conclusions of the phase-separated nature of YAP foci on the observations that the foci are round and that they are able to fuse with each other. In our case, the foci were mostly immobile and often exhibited stable elongated shapes and interconnecting bridges between foci. Notably, this does not necessarily mean that OSFs are not phase separated droplets, however, if they are, they are being deformed by their surroundings and are trapped between other entities, indicating that they are taking up most of the available volume of liquid.

We also envisioned an alternative, and, possibly, somewhat simpler explanation. This was based on the fact that increased environmental osmolarity exerts pressure onto the cell, which causes water efflux from the cytoplasm and a concomitant decrease in cellular and cytoplasmic volume. Since OSF proteins are primarily cytosolic, but most of the cytosolic proteins we observed did not form OSFs, we posited that the cytosol was structured in a way that its components were differentially affected by hyperosmotic shock. Specifically, it could be that some areas of the cytosol easily lose water (we term these areas “liquid”) and some do so less readily (“solid”), possibly due to structural rigidity. Then such areas could have different effects on cytosolic proteins, if some proteins could easily travel between the liquid and solid areas, while other could not.

During hyperosmosis water should preferentially escape from liquid parts of the cell but not from its more “solid” areas. Thus, liquid areas of the cytoplasm should decrease in volume, forcing solid areas, as well as some organelles, to come into closer contact, thus creating pockets of liquid (Figure 8). While all of the cytosolic components increase in concentration due to reduction of cellular volume, the increase in the concentration of components trapped in the liquid network will be much more drastic, since they are unable to access the considerable volume represented by the “solid” areas. So, with GFP labeling, this would make these pockets of concentrated liquid small and bright (microscopically observable as OSFs). This idea is compatible with the idea of LLPS, i.e. the phase separated droplets could be forming in the concentrated pockets of liquid, however, a simpler explanation is that the OSFs may in fact be the liquid pockets themselves.

**Figure 8.**
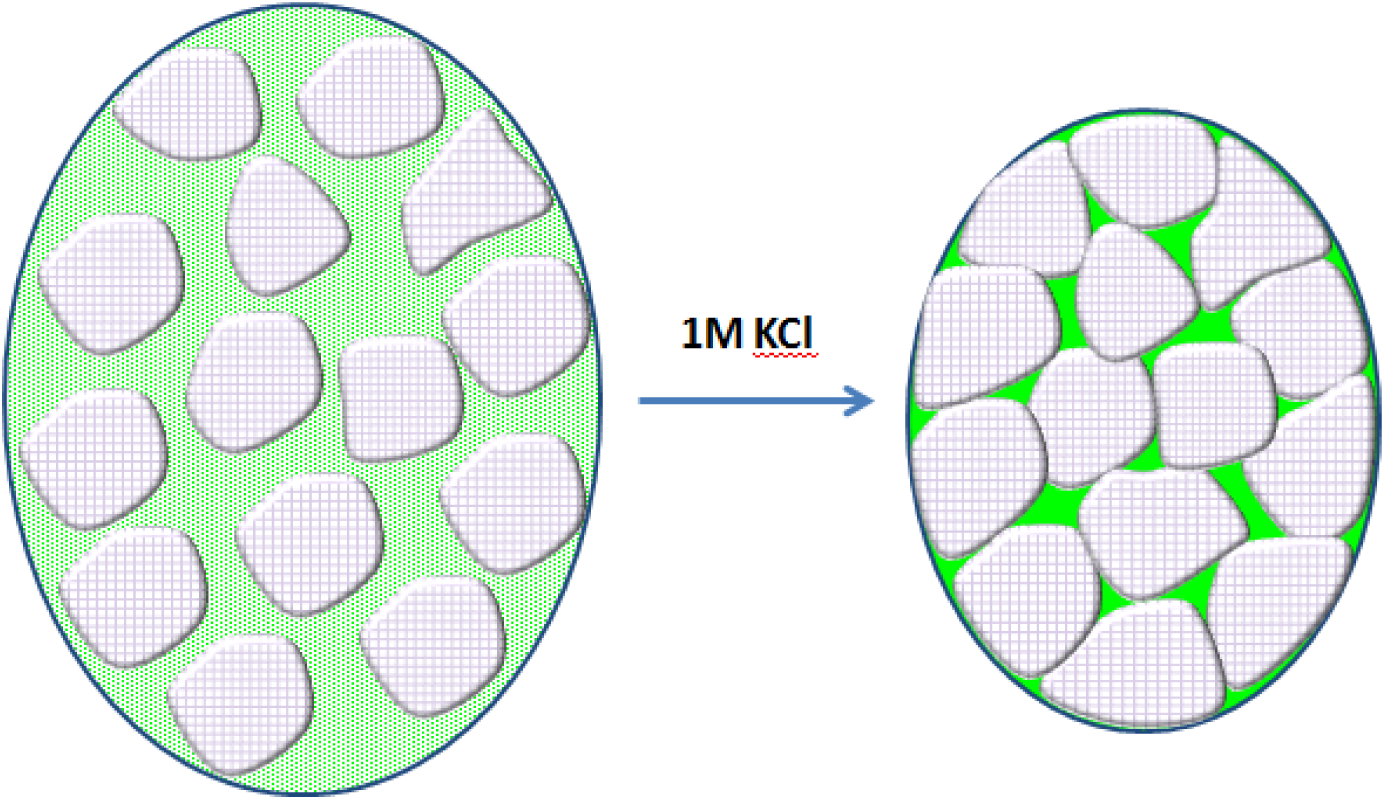
Schematic representation of the proposed mechanism of OSF emergence during hyperosmotic shock.

Notably, fusion of foci, which is a commonly accepted indicator of liquid-liquid phase separated nature of foci, is also likely in the suggested model, because shifting of “solid” entities during or after water efflux should result in an effect that could look like fusion, even though it actually may be liquid network rearrangement.

Rapid formation and disappearance are also inherent features of this model, since no processes other than change of cytoplasmic volume are required. The morphology of OSFs revealed by high-resolution microscopy is also consistent with this model, since pockets of liquid formed due to bunching together of solid compartments would be expected to look like the interconnected OSF shapes observed by SIM. Finally, screening for gene deletions and knock-downs interfering with Ssa1 OSF appearance, revealed no single deletions that significantly impacted the OSF-forming behavior of Ssa1-DDR, which is consistent with the described model. We must note however, that LLPS can also proceed very quickly under certain circumstance (Bracha et al., 2018a)

Our “beads in liquid” model implies that OSF formation should be characteristic of proteins that are trapped in the liquid phase, i.e. for proteins which cannot travel freely between the liquid and “solid” cytoplasmic compartments. What would the most likely mechanism of this trapping be? Our data suggest one likely explanation, namely, that aggregates or mutlimeric complexes formed in the liquid phase cannot easily enter the solid areas. We suppose that this might be due to a rigid sponge-like nature of the solid areas, where the pores of the sponge are large enough to admit monomeric or oligomeric proteins, but cannot accommodate large protein assemblies. The role of protein complex size is confirmed by the role of protein aggregation in OSF formation, which was directly demonstrated for PrDSup35-GFP. In a monomeric state this protein does not participate in OSFs, probably because it can travel freely between liquid and solid areas. The same behavior is observed for the monomeric fluorescent protein Dendra2. However, PrDSup35-GFP prion polymers should not travel between compartments so freely and, therefore, are likely to be accumulated in a liquid phase of cytoplasm and this is why upon hyperosmotic shock they are collected into OSFs.

A similar consideration can be applied to involvement of chaperones, such as Ssa1, into OSFs. We observed that growth on medium with glycerol as the sole carbon source, a condition which causes increased protein carbonylation compared to growth on glucose (Vasylkovska et al., 2015), as well as mild heat and cold stress in cells grown on glucose as the sole carbon source promoted OSF formation by Ssa1, suggesting that the chaperone bound damaged proteins and thus formed large complexes, which resulted in their entrapment in the liquid phase of the cytoplasm.

Notably, most (at least 16 out of 19) of the identified proteins capable of forming OSFs, have also been shown to be components of multi-protein complexes, i.e. P-body proteins (Mugridge et al., 2018), Tps1, a component of the 800 kDa trehalose-6-phosphate synthase complex (Londesborough and Vuorio, 1991), Fas2, a component of the 2.6 MDa fatty acid synthase complex (Lomakin et al., 2007), Gln1, which forms a dodecamer nanotube-like assembly (He et al., 2009) and Ape1, forming a homododecamer and higher order complexes (Kim et al., 1997).

Our model “beads in liquid” model indicates that concentrated pockets of liquid should form under any of the conditions tested, as they are a fundamental physical response of the cytosol to hyperosmotic shock, but the presence of various proteins in OSFs would depend on whether they are involved in large aggregates located in the liquid phase under given conditions. Indeed, we observed that conditions that prevented OSF participation for Ssa1 (growth on YPD at 30°C) still allowed OSF participation for Hsp42 (Figure 1B).

Thus, if the proposed model of the cytoplasmic architecture is correct, the cytoplasm can be considered as a mixture of liquid and solid areas. The general view on the structure of the cytosol ranges from a simple solution, to a crowded liquid, to a gel, and recent data also report on the ability of the cytoplasm as a whole to transition between liquid (characterized by rapid diffusion) and glass-like states (Munder et al., 2016; Parry et al., 2014). Thus our model provides a clear mechanistic way of how all of these concepts can be united. Furthermore, the suggested hypothesis is in line with observations indicating that the cytoplasm is not homogeneous and contains areas with different diffusion properties (Luby-Phelps, 1999). Indeed, recent work, which used fluorescent correlation spectroscopy to construct diffusion maps of several cytoplasmic proteins, showed that the cytoplasm is not uniform in terms of protein diffusion rates (Ranjit et al., 2014). Also it has been shown that the cytoplasm contains areas which were less accessible to 17 nm particles as compared to 1 nm particles (Luby-Phelps, 1999), proving the existence of structures that exclude large particles. Similar observation were also made more recently (Feric et al., 2016). These works were performed using cells of higher eukaryotes, and in most cases the observed effects were attributed to structures formed by the actin cytoskeleton (Feric et al., 2016; Guo et al., 2014). However, OSF formation was not affected by disassembling the actin cytoskeleton with latruncullin A, or by benomyl, a tubulin depolymerizing agent (data not shown). Thus it is possible that various types of cytoplasmic structuring can exist, and furthermore, since yeast and other microorganisms differ in the structure of their cytoskeletons from higher eukaryotes and possess an exoskeleton in the form of their cell wall, they may accomplish the same heterogeneity of the cytoplasm *via* means other than their cytoskeleton.

It is also noteworthy that according to our data most proteins do not form OSFs. This can be because these proteins are monomeric or can form only small oligomers, which enable such proteins to travel between the putative liquid and solid phases freely or, alternatively, these proteins can preferentially inhabit “solid” compartments, the nature of which is currently unclear, but should be a very promising area of investigation.

The biological significance of the proposed cytoplasmic architecture is expected to be profound. The “beads in liquid” organization of the cytosol would provide two types of environments in which cellular processes could take place. The liquid environment should allow rapid diffusion of cytoplasmic components, which, for example, must be relevant to signaling processes in which a signal molecule must travel a considerable distance through the cell. In contrast, processes that involve repeating transient interactions between partner proteins, or sequential reactions of a substrate with a set of enzymes, would benefit from slow rates of diffusion, as described in (Guigas and Weiss, 2008). This would make the network of liquid cytoplasm somewhat analogous to the circulatory systems of higher organisms, while the solid cytoplasmic areas and organelles would be more akin to stationary tissues. Also, since our data indicate that aggregates of damaged proteins are localized to the liquid phase, this may help protect various proteins in the solid cytoplasmic compartments from pathological interactions. Lastly, presence of solid compartments, which experience changes of concentrations of certain components during osmotic shock much more slowly than the liquid components, could allow buffering of the dangerous effects of osmotic shock, shielding sensitive components. Further studies are necessary to support the suggested “beads in liquid” model of cytosolic architecture, determine its relevance for processes which take place in the cytoplasm, as well as identify the nature of the solid compartments. Importantly, our discovery provides a model in which all of these directions can be explored.

## Acknowledgements

We thank Dr. Daniel Kaganovich, Triana Amen and other members of the Kaganovich lab, as well as Dr. Vadim Gladyshev and Dr. Sergey Dmitriev for the use of materials, equipment and fruitful discussions. This work was primarily funded by the grant from the Russian Science Foundation # 17-14-01092 and the Ministry of Science and Higher Education of the Russian Federation. Experiments involving Structured Illumination Microscopy were performed with partial support of the Moscow State University development program (PNR 5.13) and Russian Federation grant 14.W03.31.0012. Work involving construction of the SGA-array and real-time imaging was, in part, supported by a Short-Term Fellowship to AIA from EMBO. High-throughput microscopy was partially supported by the Ministry of Education and Science of the Russian Federation (Agreement No. 02.A03.21.0003 dated August 28, 2013).

## Materials and methods

### Yeast strains and cultivation conditions

Most of the experiments in this work used cells derived from the BY4741 strain (MATa *his3-1 leu2-0 met15-0 ura3-0*), except for the experiments concerning OSFs formed by amyloidogenic proteins, which were performed in 74D-694 (MATa *ade1-14, trp1-289, his3Δ-200, ura3-52, leu2-3,112*) (ref) and deletion screening, which was performed on hybrids between BY4741 strains from the Yeast Deletion (Giaever *et al*., 2002) and Yeast DAmP collections (Breslow *et al*., 2008) collection with a SGA query strain Y5563 (MATα *can1Δ*::MFA1pr-HIS3 *lyp1Δ ura3Δ0 leu2Δ0 his3Δ1 met15Δ0*) (Tong and Boone, 2006). The media used were YPD (Yeast extract 1%, Peptone 2%, Glucose 2%) and SC-Gly (Yeast extract 1%, Peptone 2%, 2.5% glycerol), and SC-D (Yeast nitrogen base – 0.17 g/l, ammonium sulfate – 5 g/l, glucose – 2%, casamino acids – 0.5%, tryptophan – 75 mg/l, uracil – 75 mg/l, adenine – 19 mg/l) and SC-Gly (same, except with 2.5% glycerol instead of glucose). When necessary, solid medium was prepared by including 2% agar. Unless noted otherwise the cells were grown to mid-log phase at a temperature of 30°C.

The *Ogataea parapolymorpha* strain DL1-L (*leu2*), derived from DL-1 (ATCC 26012) was modified to produce Hsp70 tagged with tagGFP. We chose the closest homologue of *S. cerevisiae* Ssa1 (NCBI Ref XP_013937290). A codon-optimized tagGFP coding sequence (ordered from Biomatic, Canada) (Supplemental Materials) possessing SalI prior to the the tagGFP ORF was cloned between Asp718 and BglII sites of pAM773 (Agaphonov, 2017). Then, inversed recombination arms directing plasmid integration into the *O. parapolymorpha SSA1* locus were inserted between SalI and EcoRV sites of the resulting plasmid. The fragment with the inversed recombination arms was obtained by PCR using primers DL_SSA_U2 and DL_SSA_L_Xho (see Table 2 for all primer sequences) and SalI-digested and self-ligated *O. parapolymorpha* genomic DNA. The PCR product was cleaved at one side with XhoI to allow ligation with SalI-generated cohesive end of the vector. Prior to yeast transformation the obtained plasmid pAM783 was digested with SalI to obtain the cassette for replacement of the wild-type SSA for SSA-tagGFP fusion according to scheme described previously (Agaphonov et al., 2009).

**Table 2.**
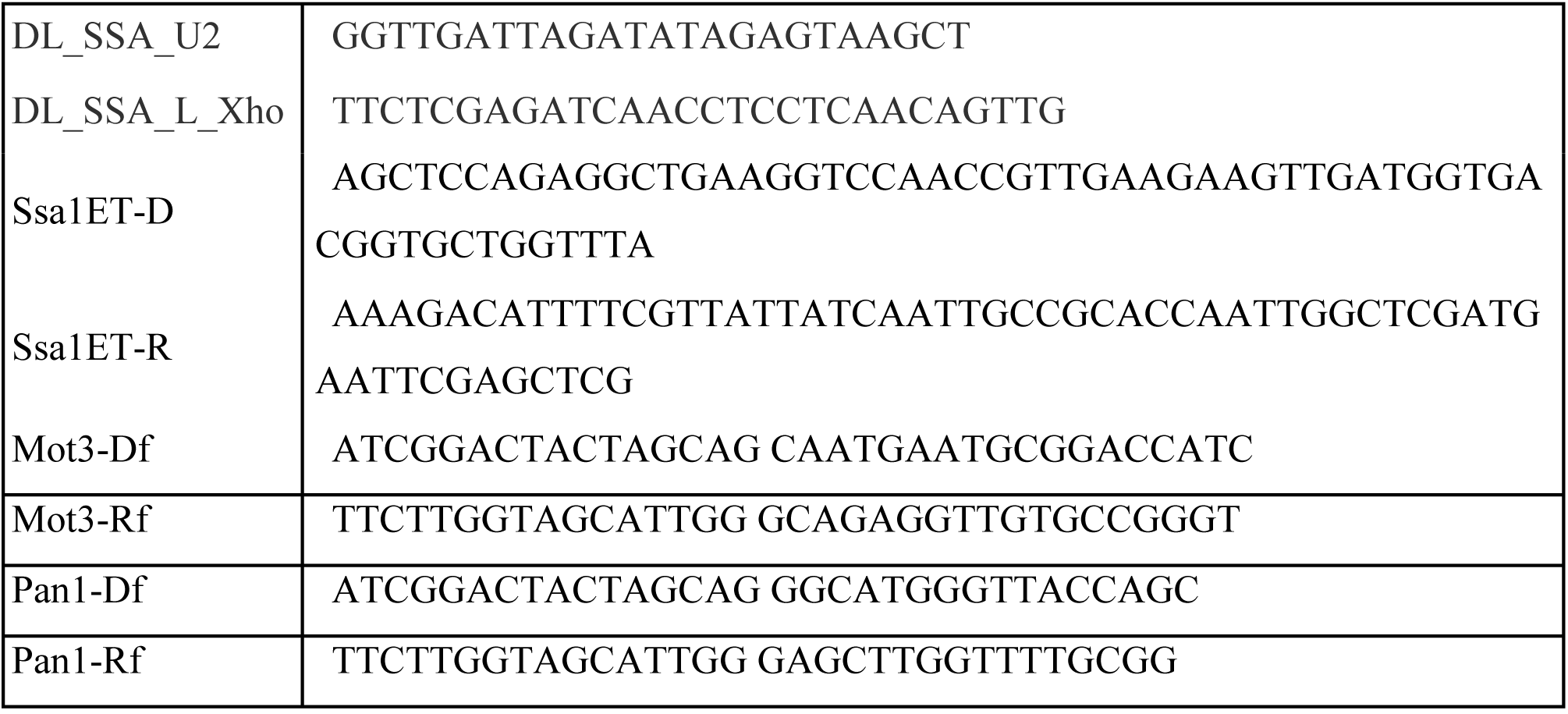
Primers used in this work.

**Table 3.** Plasmids used in this work.

|  |  |  |
| --- | --- | --- |
| pAM783 | Integrative plasmid for endogenous tagging of O. parapolyomorpha Hsp70 with tagGFP | This work |
| pAM781D | Integrative plasmid for replacement of GFP:: HIS3 module in the GFP Fusion collection with tagRFP::URA3 | This work |
| 127-DDR-hph | Template plasmid for endogenous tagging of proteins with DDR2 (HygR selection marker) | Gift from D. Kaganovich |
| pGal-HO | Autonomous plasmid for mating-type switching, *LEU2* and *URA3* auxotrophic markers, galactose-inducible promoter | unknown source |
| pYES2-Sup35NMH-GFP | Autonomous multicopy plasmid for expression of the prion domain of Sup35 tagged with GFP, galactose-inducible promoter | This work |
| pYES2-PrDMot3-GFP | Autonomous multicopy plasmid for expression of the prion domain of Mot3 tagged with GFP, galactose-inducible promoter | This work |
| pYES2-PrDPan1-GFP | Autonomous multicopy plasmid for expression of the amyloidogenic domain of Pan1 tagged with GFP, galactose-inducible promoter | This work |

Strains producing tagRFP-fusions were constructed from appropriate GFP-tagged strains by using a universal plasmid that replaces the GFP::*HIS3* cassete from the Yeast GFP collection with a tagRFP::*URA3* cassete. The plasmid was constructed by modifying the plasmid pFA6a– GFP(S65T)–His3MX, which contains the *Schizosaccharomyces pombe HIS5* gene capable of complementing the *S. cerevisiae his3* mutation (Longtine et al., 1998). The SmaI-BsrGI fragment was replaced with the g-block tagRFP_Sc (IDT, USA) possessing a codon-optimized tagRFP sequence (See supplement for sequences of synthetic genes). Then the NcoI-ScaI fragment bearing the *HIS5 S. pombe* ORF in the resulting plasmid was replaced with PciI-DraI fragment of the *O. polymorpha URA3* locus. The resulting plasmid designated as pAM781D (See supplement for plasmid maps) was digested with SalI-ClaI to obtain cassette bearing tagRFP with *URA3* selectable marker flanked with sequences homologous to flanking regions of the GFP(S65T)–*His3MX* encoding module.

Strains for colocalization studies were obtained by switching the mating type of the obtained tagRFP-fusion strain with pGal-HO, a plasmid encoding the HO-endonuclease gene under control of an inducible *GAL1* promoter, verifying mating type change, and crossing it with appropriate GFP-fusion strains. Diploids were selected on SC–His,-Ura medium. To obtain haploids, the diploids were sporulated in liquid 1% potassium acetate for 5 days and then, after verifying formation of asci, suspended in distilled water mixed 1:1 with ether (Dawes and Hardie, 1974) and incubated for 30 minutes. These suspensions were plated onto SC-D medium lacking His and Ura to select for segregants encoding both fusion proteins as verified by microscopy and being haploid as verified by assaying mating type.

The strains producing Ssa1-DDR were created by PCR-based endogenous tagging of BY4741 using PCR products obtained using primers Ssa1ET-D and Ssa1ET-R with the plasmid 127-DDR-hph used as a templates. The plasmid was a kind gifts from Dr. Daniel Kaganovich and papers describing them are currently in preparation (maps are included in the Supplemental Materials). The SGA query strain producing Ssa1-DDR was constructed using an identical procedure.

Plasmids encoding amyloidogenic domains tagged with GFP were constructed using the plasmid pYES2-Sup35NMH-GFP, which was created by inserting the BglII-XbaI fragment encoding Sup35(1-239)-HHHHHHPVAT-eGFP into the BamHI and XbaI sites of pYES2 (Invitrogen). Genomic DNA fragments encoding first 297 amino acid residues of Mot3 and first 223 residues of Pan1 were amplified with primers Mot3-DF, Mot3-Rf, Pan1-Df and Pan1-Rf (see Table 2), and inserted into PvuII/BalI treated pYES2-Sup35NMH-GFP using quick-fusion cloning kit (Bimake) to replace fragment encoding Sup35 residues 1-154.

### Sample preparation for microscopy

To visualize OSFs, cells were spotted onto a 2% agar pad based on SC medium containing 1M KCL (Alexandrov and Dergalev, 2019). Identical pads without KCl were used as a control. Time-lapse videos were acquired using glass bottom plates treated with concanavalin A in order to prevent cell movement.

### Confocal microscopy to study protein colocalization and for real-time imaging

Images were acquired using a dual point-scanning Nikon A1R-si microscope equipped with a PInano Piezo stage (MCL), using a 60x PlanApo VC oil objective Numerical aperture (NA)=1.40.

### High throughput microscopy

Cells were grown in deep-well 96-well plates in YP-Gly medium, washed with YP-Gly and moved to 384 glass-bottom plates (CellVis, P384-1.5H-N, USA). Each sample was placed into two adjacent wells and after loading, KCl was added up to a final concentration of 1M to one of the wells in a pair.

Imaging was performed using an ImageXpress Micro XL (Molecular Devices, USA) high-throughput microscope equipped with a 100x LWD NA=0.9. Three images were collected for each well. Images were viewed manually, and all strains exhibiting shock-induced protein relocalization were reassessed using a manual Zeiss AxioSkop 40 microscope, 100x oil immersion objective, NA=1.2.

### Structural Illumination Microscopy

Live imaging was performed on an N-SIM superresolution system (Nikon, Japan) equipped with 100x Plan Apo TIRF lens (NA=1.49) and iXon 897 EM-CCD camera (effective pixel size 63 nm) (Andor, Ireland) in 3D-SIM mode (excitation laser line 488 nm, 120 nm Z-steps) under control of NIS-Elements 4.6 software. Raw image stacks (3 grating angles x 5 phase shifts) were analyzed for image quality with SIMcheck module of ImageJ software and processed using SIM module of NIS-Elements using parameters selected on the basis of Fourier transform analysis. Reconstructed stacks were further deconvolved using the Richardson-Lucy algorithm built into NIS-Elements.

### Cytoplasmic shrinkage measurements

Cytoplasmic shrinkage analyses used images obtained from a timelapse of OSF formation and dissolution obtained on a confocal microscope (see Confocal microscopy). Using ImageJ software, single cells from the timelapse-series were transformed into shapes using the Threshold function and then the areas of these shapes were calculated using the magic wand feature. OSF formation was quantified and graphed by selecting a region of interest which contained a single OSF and the relative signal intensity in this region of interest at different timepoints was plotted onto a graph together with the change in cell area.

### Deletion screening

The query strain for the Synthetic Genetic Array producing Ssa1-DDR was used to obtain an SGA-array (Tong and Boone, 2006) using the Yeast deletion collection and the Yeast DaMP (Breslow et al., 2008) collection in 384 well format. Briefly, this involved mating, sporulation, and subsequent selection steps for obtaining haploid progeny of a specified mating type and containing both the gene deletions from the collection and the marker of interest (SSA1-DDR::HygR). The arrayed strains were then grown overnight on YPD plates and then inoculated into 384-well microscopy plates containing SC medium with glucose as a sole carbon source and 1M KCl. The plates were then imaged on a high-throughput microscope, as detailed in the High-Throughput microscopy section. All the images were then inspected manually in order to detect candidates. Strains which were judged to be defective in OSF formation were reassessed using a manual Zeiss AxioSkop 40 microscope, 100x oil immersion objective, NA=1.2.

